# Main protease mutants of SARS-CoV-2 variants remain susceptible to nirmatrelvir (PF-07321332)

**DOI:** 10.1101/2021.11.28.470226

**Authors:** Sven Ullrich, Kasuni B. Ekanayake, Gottfried Otting, Christoph Nitsche

## Abstract

The COVID-19 pandemic continues to be a public health threat. Multiple mutations in the spike protein of emerging variants of SARS-CoV-2 appear to impact on the effectiveness of available vaccines. Specific antiviral agents are keenly anticipated but their efficacy may also be compromised in emerging variants. One of the most attractive coronaviral drug targets is the main protease (M^pro^). A promising M^pro^ inhibitor of clinical relevance is the peptidomimetic nirmatrelvir (PF-07321332). We expressed M^pro^ of six SARS-CoV-2 lineages (C.37 Lambda, B.1.1.318, B.1.2, B.1.351 Beta, B.1.1.529 Omicron, P.2 Zeta), each of which carries a strongly prevalent missense mutation (G15S, T21I, L89F, K90R, P132H, L205V). Enzyme kinetics showed that these M^pro^ variants are similarly catalytically competent as the wildtype. We show that nirmatrelvir has similar potency against the variants as against the wildtype. Our *in vitro* data suggest that the efficacy of the specific M^pro^ inhibitor nirmatrelvir is not compromised in current COVID-19 variants.

**Graphical abstract:** 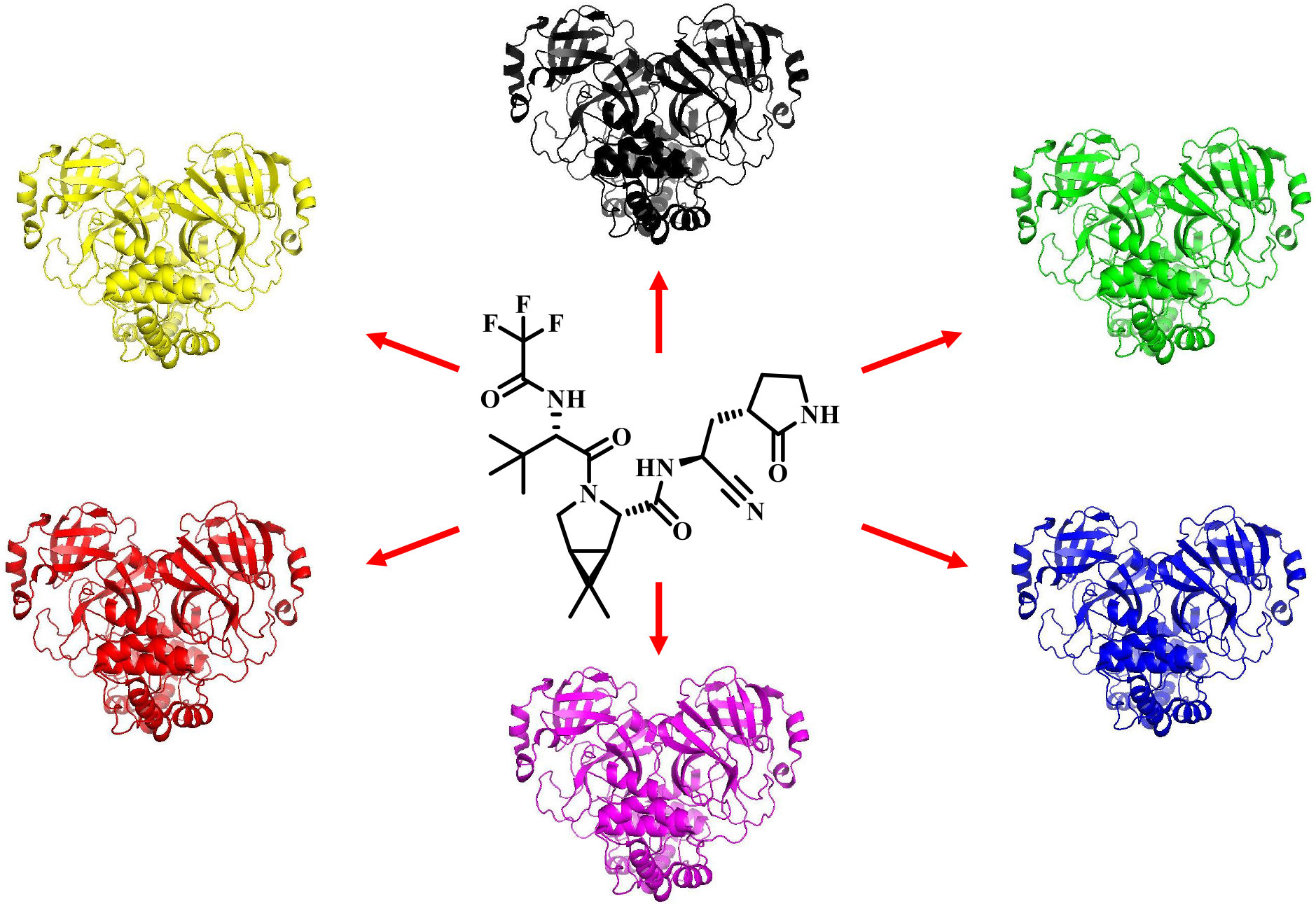

---

Since its emergence in late 2019,^1^ COVID-19 has significantly impacted on societies worldwide.^2^ More than 5 million deaths have been attributed to COVID-19, with the number of confirmed SARS-CoV-2 infections surpassing 275 million.^3^ The outbreak of SARS-CoV-2 prompted multiple successful vaccine development campaigns.^4^ Currently approved vaccines, such as viral vector or mRNA vaccines, successfully limited the pandemic’s impact on global health.^5, 6^ Most COVID-19 vaccines function by stimulating an immune response against the SARS-CoV-2 spike protein (S)^7-9^ but, as the spike gene has gathered pronounced genetic variability,^10, 11^ it is a concern if the effectiveness of existing vaccines are affected by variants of SARS-CoV-2.^5, 6, 10, 12^ At the time of writing, the World Health Organization (WHO) lists five variants of concern (VOC; Alpha, Beta, Gamma, Delta, Omicron) and two variants of interest (VOI; Lambda, Mu).^13^ A possible reformulation of the vaccines adjusted to currently circulating lineages of SARS-CoV-2 is being investigated.^14-16^ It is clear that the deployment of vaccines remains the best public health measure to control the spread of SARS-CoV-2 and the severe health effects of COVID-19.^17, 18^

Complementary to preventive vaccines, antiviral drugs are urgently needed to combat COVID-19.^19^ Since the discovery of SARS-CoV-1 in 2003,^20^ several coronaviral drug targets have been identified,^21^ including the RNA-dependent RNA polymerase (RdRp, nsp12),^22^ the helicase (nsp13),^23^ the papain-like protease (PL^pro^, part of nsp3),^24^ and the main protease (M^pro^, 3CL^pro^, nsp5).^25^ Despite this, treatment options for COVID-19 are limited. Only very recently, the orally active drugs molnupiravir (MK-4482, EIDD-2801, Lagevrio™) and nirmatrelvir (PF-07321332, Paxlovid™ as combination drug with ritonavir as booster) were approved for emergency use in the United Kingdom and the United States in November and December 2021. Molnupiravir targets RdRp by acting as a nucleoside analogue prodrug, but was originally developed against different RNA viruses.^26^ Nirmatrelvir is an orally available peptidomimetic targeting Mpro, employing a nitrile warhead to covalently bind the catalytic cysteine residue in the active site of the protease (**Figure 2a**).^27^

SARS-CoV-2 M^pro^ is a homodimeric cysteine protease, which processes the majority of the viral polyproteins pp1a and pp1ab encoded by the ORF1a/b gene.^25, 28^ Inhibition of M^pro^ thus ultimately hinders the assembly of the replication and transcription complexes (RTCs).^25, 29^ The protease has a distinct recognition motif, with – in the Schechter-Berger notation^30^ – preference for leucine in P_2_ and especially strong preference for glutamine in P_1_.^25, 31^ Human host proteases have different substrate specificities and it is therefore anticipated that selective inhibitors have limited off-target effects.^25^

Previous research on SARS-CoV-1 M^pro^ (which is 96% identical in amino acid sequence to SARS-CoV-2 M^pro^)^25^ demonstrated that missense point mutations can influence protease activity. Mutants have been identified with slightly enhanced (Ser284, Thr285, Ile286)^32, 33^ and slightly or severely reduced catalytic activity (Gly11, Asn28, Ser139, Phe140, Glu166, Asn214, Arg298).^32, 34-38^ Specifically the R298A mutation has become a tool to study the protease in its monomeric form, since it inactivates the protease by disrupting the M^pro^ dimer.^35^ The present study assesses the M^pro^ mutants of emerging SARS-CoV-2 lineages. We analysed the most widespread amino acid substitutions in SARS-CoV-2 M^pro^, characterized them by enzyme kinetics and assessed their susceptibility to inhibition by nirmatrelvir.

Utilizing the Outbreak.info database by Scripps Research,^39^ which partially operates with data provided by the GISAID Initiative,^40^ we performed an analysis of the genomes of SARS-CoV-2 lineages, including the VOC and VOI. The WIV04 sequence (EPI_ISL_402124)^41^ acted as wildtype (WT) reference genome. Lineage comparison^42^ of VOCs and VOIs revealed two missense mutations in the M^pro^ section of the ORF1a/b gene with >20% frequency of occurrance. The mutations are G15S, which is >85% prevalent^43^ in the Lambda VOI (or C.37, using PANGO nomenclature)^44^, K90R, which is >95% prevalent^45^ in the Beta VOC (B.1.351) and P132H, which is >95% prevalent^46^ in the Omicron VOC (B.1.1.529). The Delta VOC (B.1.617.2), which was the dominant lineage for most of the second half of 2021,^47, 48^ did not display any particularly prevalent (>20%)^42^ missense mutations within the M^pro^ part of ORF1a/b, implying that its M^pro^ is identical to that of the WT. We additionally chose to investigate three additional abundant M^pro^ mutations to cover a larger variety of lineages: T21I, which is >90% prevalent^49^ in B.1.1.318, a WHO variant formerly under monitoring (VUM),^13^ L89F, which is >95% prevalent^50^ in the B.1.2 lineage, and L205V, which is >95% prevalent^51^ in the former VOI Zeta (P.2) (**Figure 1b**). Hence, we selected the six mutations G15S, T21I, L89F, K90R, P132H and L205V for further investigations.

**Figure 1.**
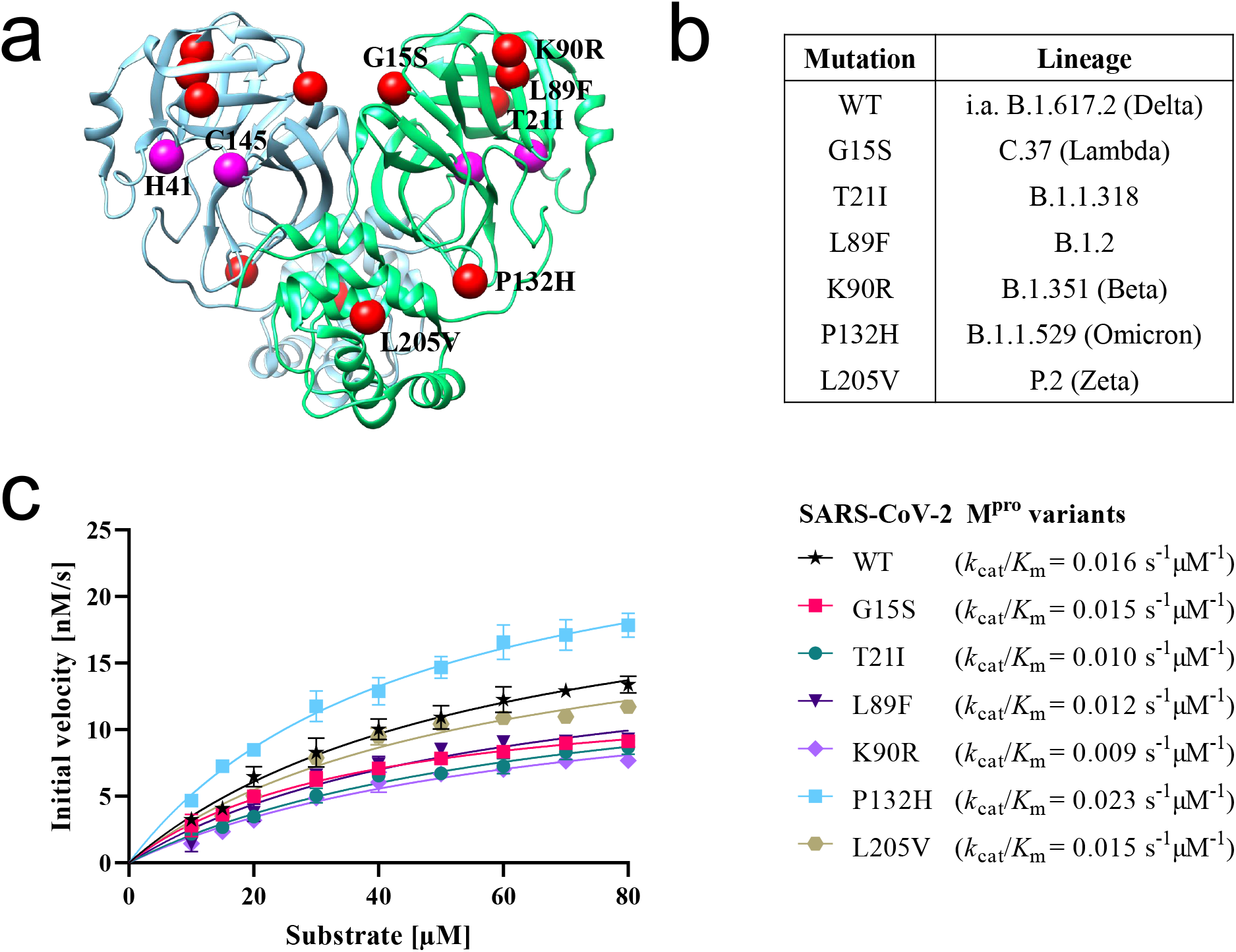
Comparison of M^pro^ mutations and proteolytic activities. (a) X-ray structure of SARS-CoV-2 M^pro^ (PDB: 6LU7)^52^ indicating the location of mutations (red) and the catalytic dyad (purple) in the two protomers (cyan, green). (b) List of prevalent M^pro^ mutations and their corresponding SARS-CoV-2 lineage. (c) Michaelis-Menten kinetics of SARS-CoV-2 M^pro^ variants specifying their catalytic efficiency (*k*_cat_/*K*_m_).

X-ray crystal structures of WT SARS-CoV-2 M^pro^ (e.g. PDB: 6LU7)^52^ indicate that the residues G15, T21, K90 and P132 are solvent-exposed, while the hydrophobic residues L89 and L205 are buried within the protease. Except for T21I and P132H, the mutations introduce no major changes in the chemical character of the side-chains, as indicated by low – or in the case of T21I and P132H moderate – values of Miyata’s distances.^53^ The mutations G15S, T21I, L89F and K90R are located in domain I, while the mutations P132H and L205V are in domains II and III, respectively (**Figure 1a**).^25^

WT SARS-CoV-2 M^pro^ and the mutants G15S, T21I, L89F, K90R, P132H and L205V were expressed in *E. coli* and purified. An established Förster resonance electron transfer (FRET) *in vitro* assay of M^pro^ activity^54^ was employed to determine initial velocities of the proteolytic activity at various substrate concentrations. The data confirmed that all mutants are enzymatically active, which was expected^25, 55^ as a dysfunctional M^pro^ would prevent replication of SARS-CoV-2. The seven M^pro^ variants exhibited turnover numbers (*k*_cat_) between 0.54 and 1.03 s^-1^, and Michaelis constants (*K*_m_) ranging from 37 to 67 µM (**Table S1**). The catalytic efficiencies (*k*_cat_/*K*_m_) calculated for the mutants (0.009 to 0.023 s^-1^µM^-1^) are similar to that of WT M^pro^ (0.016 s^-1^µM^-1^), confirming that all M^pro^ variants are equally competent with regard to their proteolytic activities (**Figure 1c, Table S1**).

Following the kinetic analysis of the SARS-CoV-2 M^pro^ variants, the clinical candidate nirmatrelvir (also known as PF-07321332 ; **Figure 2a**) was used to assess the potential impact of M^pro^ mutations on the drug’s efficacy. The inhibition constant (*K*_i_) of nirmatrelvir against SARS-CoV-2 WT M^pro^ has been reported to be 3.1 nM.^27^ Our FRET assay confirmed that nirmatrelvir inhibits the activity of M^pro^ variants at nanomolar compound concentrations. Furthermore, the extent of inhibition was similar across the different protease variants, with 5 nM compound displaying inhibition below 50%, 20 nM showing inhibition over 50% and 100 nM fully inhibiting the enzymatic activity of all mutants and the WT (**Figure 2b**).

**Figure 2.**
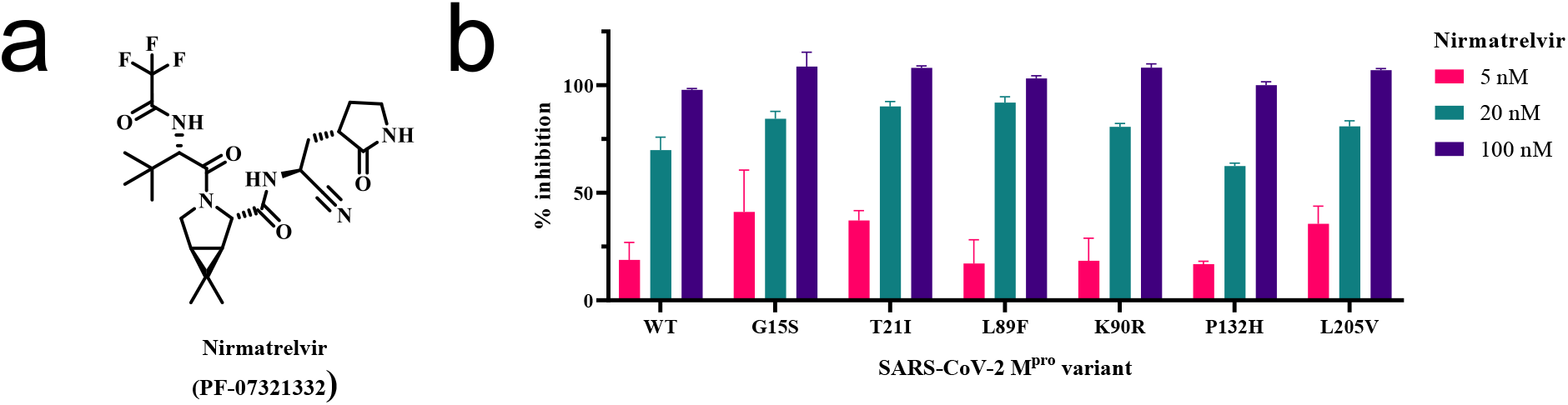
Inhibition of SARS-CoV-2 M^pro^ variants. (a) Chemical structure of the inhibitor nirmatrelvir^27^ (b) *In vitro* inhibition of SARS-CoV-2 M^pro^ variants by nirmatrelvir.

In summary, we identified the currently most prevalent M^pro^ variants (G15S, T21I, L89F, K90R, P132H, L205V) in different lineages of SARS-CoV-2 (C.37 Lambda, B.1.1.318, B.1.2, B.1.351 Beta, B.1.1.529 Omicron, P.2 Zeta) and found that in a biochemical assay they are similarly catalytically competent as the wildtype. In addition, we confirmed that nirmatrelvir maintains effective inhibition of all these M^pro^ variants *in vitro*. This gives hope that nirmatrelvir and potentially other specific SARS-CoV-2 M^pro^ inhibitors would presently not be affected negatively by these virus variants. It must be noted, however, that widespread use of M^pro^ inhibitors may challenge SARS-CoV-2 to develop M^pro^ mutations that overcome these inhibitors, as previously experienced for, e.g., HIV protease inhibitors.^56^ Despite these challenges, protease inhibitors have revolutionized antiviral treatment for viral infectious diseases, including HIV and HCV.^57^ It can thus be expected that M^pro^ inhibitors will have a similar impact on the future development of the COVID-19 pandemic.

## Acknowledgements

C.N. thanks the Australian Research Council (ARC) for a Discovery Early Career Research Award (DE190100015) and Discovery Project funding (DP200100348). G.O. thanks the ARC for a Laureate Fellowship (FL170100019) and acknowledges support by the ARC Centre of Excellence for Innovations in Peptide & Protein Science (CE200100012). This study was supported by a RAMR (MAWA) grant awarded to S.U. and C.N.

