## Supporting Information for "Main protease mutants of SARS-CoV-2 variants remain susceptible to nirmatrelvir (PF-07321332)"

**Abbreviations**

| | |
|---|---|
| CI | Confidence interval |
| CoV | Coronavirus |
| DABCYL | 4-((4-(Dimethylamino)phenyl)azo)benzoic acid |
| DTT | Dithiothreitol |
| EDANS | 5-((2-Aminoethyl)amino)naphthalene-1-sulfonic acid |
| EDTA | Ethylenediaminetetraacetic acid |
| FRET | Förster resonance electron transfer |
| HEPES | 4-(2-Hydroxyethyl)-1-piperazineethanesulfonic acid |
| IPTG | Isopropyl-β-D-thiogalactopyranoside |
| $M^{pro}$ | Main protease |
| OD | Optical density |
| PAGE | Polyacrylamide gel electrophoresis |
| SARS | Severe acute respiratory syndrome |
| SDS | Sodium dodecyl sulphate |
| UHPLC | Ultra high-performance liquid chromatography |
| WT | Wildtype |

### Plasmid construction

The SARS-CoV-2 $M^{pro}$ gene[1] was cloned in-between the *NdeI* and *XhoI* sites of the T7 vector pET-47b (+). The construct contains the $M^{pro}$ cleavage-site (SAVLQ↓SGFRK; arrow indicating the cleavage site) at the N-terminus and a modified PreScission cleavage site (SGVTFQ↓GP) followed by a $His_6$-tag at the C-terminus. Point mutagenesis to incorporate an amino acid of choice at selected sites was done using overlapping primers. All plasmid constructions and mutagenesis were conducted with a RQ-SLIC and QuikChange protocol using mutant T4 DNA polymerase.[2]

### Protein expression

To produce WT $M^{pro}$ and mutants, *E. coli* BL21 DE3 cells were transformed with the respective pET-47b (+)-$M^{pro}$ plasmid and the cells were grown at 37 °C in LB medium containing 50 mg/L kanamycin. Overnight cultures were inoculated into fresh terrific broth (TB) medium (1:100 dilution) supplemented with 50 mg/L kanamycin. The cells were grown at 37 °C to an $OD_{600}$ of 0.6 to 1.0. Expression was initiated by the addition of IPTG to a final concentration of 1 mM. Protein expression was conducted by incubation at room temperature overnight.

### Protein purification

Cells were harvested by centrifugation at 5,000 *g* for 15 min. Following resuspension in buffer A (50 mM Tris-HCl, pH 7.5, 300 mM NaCl), the cells were lysed using an Emulsiflex C5 (Avestin, Canada) with two passes using 10,000–15,000 psi. The cell lysates were centrifuged for 1 h at 30,000 *g*. The supernatant was loaded onto a 1 mL His GraviTrap TALON® column (Cytiva, United States). The column was washed with 20 column volumes buffer B (50 mM Tris-HCl, pH 7.5, 300 mM NaCl, 5 mM imidazole) and the protein was eluted with 5 column volumes buffer C (50 mM Tris-HCl, pH 7.5, 300 mM NaCl, 300 mM imidazole). The fractions were analysed by 12% SDS-PAGE. Following purification, the buffer was exchanged to buffer D (20 mM HEPES-KOH, pH 7.0, 150 mM NaCl, 1 mM DTT, 1 mM EDTA) using an Amicon ultrafiltration centrifugal tube (Merck Millipore, United States) with a molecular weight cut-off of 10 kDa. All samples were analysed by mass spectrometry using an Orbitrap Fusion Tribrid Mass Spectrometer (Thermo Scientific, United States) coupled with an UltiMate S4 3000 UHPLC (Thermo Scientific, United States). Protein concentrations were determined by measuring the absorbance at 280 nm, using $\varepsilon = 33{,}640\ L \cdot M^{-1} \cdot cm^{-1}$.

**FRET assay**

The SARS-CoV-2 M$^{pro}$ FRET assay employed for the kinetic evaluation and the inhibitory assessment was based on the SARS-CoV-1 M$^{pro}$ assay by Zhu et al.[3] and the SARS-CoV-2 M$^{pro}$ assay from Zhang et al.[1] Specific conditions were adapted from Ullrich et al.[4] Black 96-well polypropylene plates with U-bottom (Greiner Bio-One, Austria) were used to carry out the assay and 20 mM Tris-HCl pH 7.3, 100 mM NaCl, 1 mM EDTA, 1 mM DTT was used as buffer. The peptide DABCYL-KTSAVLQ↓SGFRKM-E(EDANS)-$NH_2$ (Mimotopes, Australia) was used as FRET substrate. The concentration of the recombinant SARS-CoV-2 M$^{pro}$ variants amounted to 25 nM. Inhibitor nirmatrelvir[5] – also known as PF-07321332 – (MedChemExpress, United States; Batch HY-138687-116180)[6] was used to assess the inhibition of the proteases. For the kinetic assessment, FRET substrate concentrations between 10 µM and 80 µM were used, and the inhibition assay was conducted with a FRET substrate concentration of 25 µM. When inhibitor was used, it was incubated with the protease for 10 min at 37 °C. The enzymatic reaction was initiated by the addition of FRET substrate and monitored at 37 °C for 5 min at $\lambda_{em}$ = 490 nm with an excitation wavelength of $\lambda_{ex}$ = 340 nm, using an Infinite 200 PRO M Plex fluorophotometer (Tecan, Switzerland). Kinetic measurements were performed in duplicate and inhibitory measurements in triplicate. 100% enzymatic activity was defined as the initial velocity of the control reactions and %-inhibition was calculated accordingly. Fluorescence intensity was converted to cleaved substrate per time unit with an EDANS calibration curve.[7] The data obtained were analyzed and visualized with Prism 9.3 (GraphPad Software, United States).

**Amino acid sequence of SARS-CoV-2 M$^{pro}$ WT**

SGFRKMAFPSGKVEGCMVQVTCGTTTLNGLWLDDVVYCPRHVICTSEDMLNPNYE
DLLIRKSNHNFLVQAGNVQLRVIGHSMQNCVLKLKVDTANPKTPKYKFVRIQPGQT
FSVLACYNGSPSGVYQCAMRPNFTIKGSFLNGSCGSVGFNIDYDCVSFCYMHHMELP
TGVHAGTDLEGNFYGPFVDRQTAQAAGTDTTITVNVLAWLYAAVINGDRWFLNRF
TTTLNDFNLVAMKYNYEPLTQDHVDILGPLSAQTGIAVLDMCASLKELLQNGMNGR
TILGSALLEDEFTPFDVVRQCSGVTFQ

**Table S1.** Michaelis-Menten parameters of SARS-CoV-2 $M^{pro}$ variants. 95% CI in brackets.

| Variant | $k_{cat}$ [$s^{-1}$] | $K_m$ [µM] | $k_{cat}/K_m$ [$s^{-1}µM^{-1}$] |
|---|---|---|---|
| WT | 0.94 (0.77–1.11) | 57 (38–76) | 0.016 (0.008–0.024) |
| G15S | 0.54 (0.49–0.59) | 37 (29–45) | 0.015 (0.010–0.020) |
| T21I | 0.64 (0.51–0.77) | 67 (43–91) | 0.010 (0.005–0.015) |
| L89F | 0.69 (0.87–0.51) | 58 (30–86) | 0.012 (0.003–0.021) |
| K90R | 0.60 (0.46–0.74) | 67 (40–94) | 0.009 (0.003–0.015) |
| P132H | 1.03 (0.90–1.16) | 45 (33–57) | 0.023 (0.014–0.032) |
| L205V | 0.82 (0.65–0.99) | 55 (34–76) | 0.015 (0.006–0.024) |

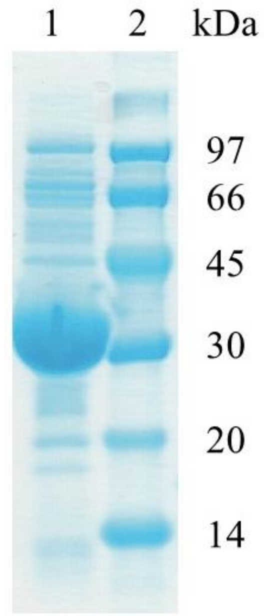


**Figure S1.** 12% SDS-PAGE images of WT SARS-CoV-2 $M^{pro}$. Lane 1 corresponds to the purified protein and lane 2 displays the protein markers.

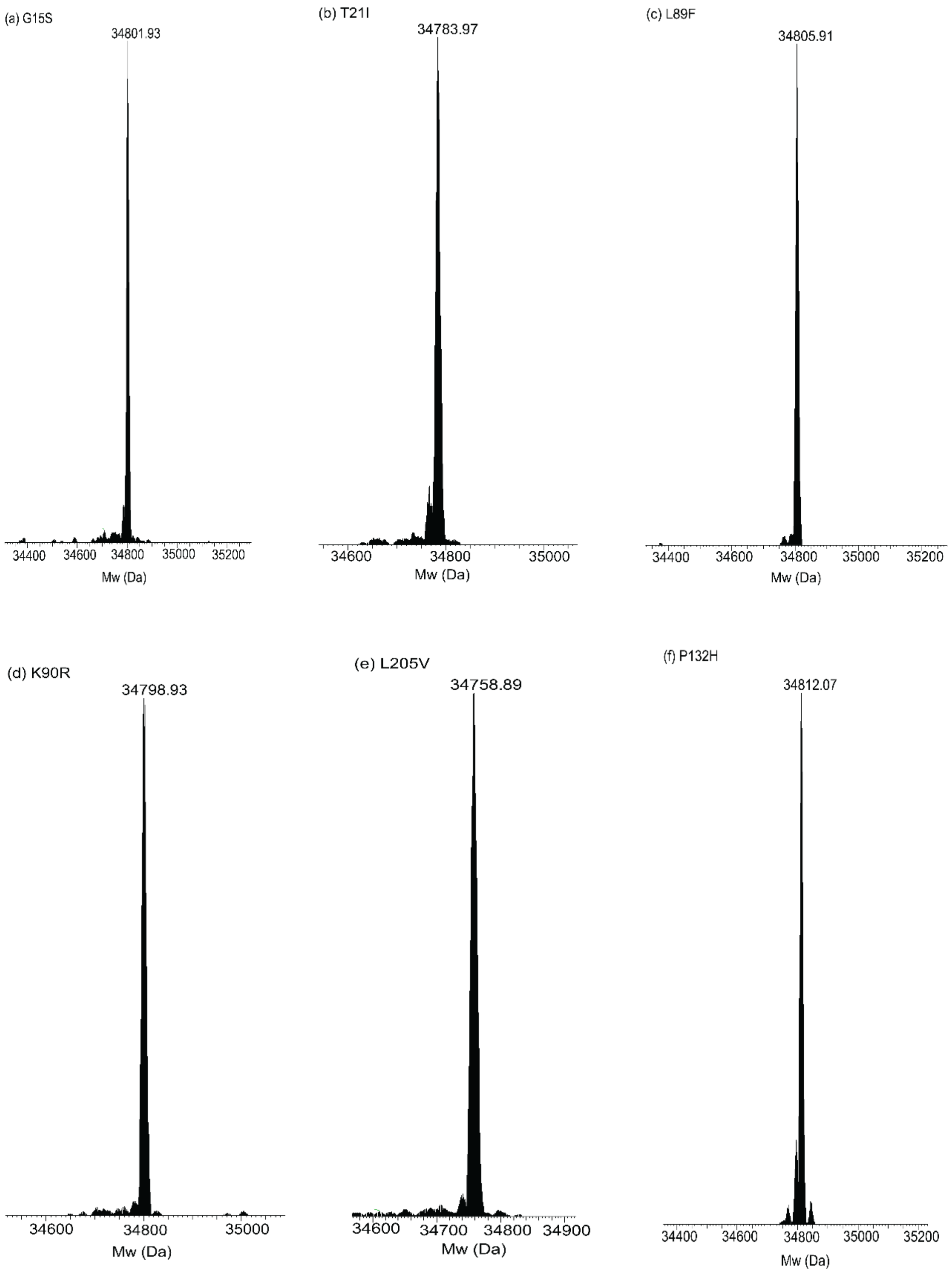


**Figure S2.** Intact protein mass spectrometric analysis of mutants of SARS-CoV-2 M$^{pro}$. Calculated molecular weights in brackets. (a) G15S (34803.77 Da). (b) T21I (34785.80 Da). (c) L89F (34807.76 Da). (d) K90R (34801.76 Da). (e) L205V (34759.72 Da). (f) P132H (34813.77 Da).
